# The role of sleep in the retention and long-term maintenance of newly formed ability self-beliefs

**DOI:** 10.64898/2026.09.14.751019

**Authors:** Julia Carbone, Christine Barner, Laura Encica, Jan Born, Susanne Diekelmann, Nora Czekalla, Alexander Schröder, Laura Müller-Pinzler, Sören Krach

## Abstract

Sleep supports the consolidation of newly acquired information, however its role in the formation of self-beliefs has not yet been examined. Here, we investigated whether sleep facilitates the retention, stabilization, revision and long-term maintenance of newly formed ability beliefs. Using a Sleep vs. Wake between-subjects design (N=54), participants completed the Learning Of Own Performance (LOOP) task, which allowed them to form new positive and negative ability self-beliefs based on mocked feedback at a first session, and to revise them when encountering contradictory feedback at a second session. Belief formation was followed by either a sleep or wake retention interval. Belief states were tested at four time points: immediately after belief formation, after the 12-h retention interval, following the revision phase and after three weeks. Participants successfully formed and revised self-beliefs using the LOOP task. Compared with wakefulness, sleep following belief formation did not affect belief retention across the 12-h interval, nor did it stabilize beliefs against their revision. However, after three weeks, participants who slept after belief formation showed a shift towards more positivity for the initially formed low ability self-beliefs. Contrary to our hypotheses, sleep did not enhance short-term retention or stabilization of newly formed self-beliefs. However, the more positively recalled ability self-belief associated with the initially low-performance condition suggest that sleep may facilitate a positive long-term shift.

## Introduction

Beliefs we hold about ourselves (i.e. self-beliefs) guide our behavior and social interactions. At the same time, self-beliefs are continuously shaped by social interactions and the feedback we receive during these interactions ^1–3^. Social feedback thus helps us evaluate how well we perform, how others perceive us, and whether we meet social expectations. However, social information is inherently complex. To build a coherent and stable sense of self, individuals must reduce this complexity by abstracting and integrating meaningful information from social interactions. Yet, it remains unclear how social feedback is abstracted during the process of self-belief formation and how it is consolidated over time.

Previous research shows that sleep supports the consolidation of newly encoded information by strengthening and integrating them into pre-existing knowledge networks for long-term storage^4,5^. In this consolidation process, sleep has also been proposed to promote the abstraction of gist-like information^6,7^, for example, the identification of invariant features in experience and reorganization into schema-like knowledge, thereby enhancing long-term memory access and generalization^8,9^. Gist abstraction may benefit from both slow wave sleep (SWS)^8,10^ as well as rapid-eye movement (REM) sleep^11–14^. In the last years, increasing evidence also indicates that sleep can strengthen newly formed beliefs, including stereotypes about gender and race^15–19^, however, it is currently not clear whether sleep likewise is related to how self-related beliefs are formed^20^.

Self-beliefs can be considered schema-like representations that are continuously updated based on social feedback^21,22^. Therefore, abstracting information from self-related feedback, for example in the context of ability, is necessary to form broader self-beliefs (i.e., depending on the psychological context, the term belief is often equated with other related psychological terms such as attitude, opinion, self-image, mindset, schema-like representation, or self-concept etc.) ^23–25^. The formation and revision of self-beliefs do not only depend on the content of the feedback but also on the a-priorly established value ^26^. Previous findings suggest that there is no whatsoever objective integration of feedback, but rather individuals are inherently biased in how they form self-related beliefs (e.g., such as a tendency to confirm their prior self-beliefs) ^2,27–29^. While some studies show that participants tend to update their beliefs more strongly after positive feedback (e.g., optimistic bias regarding one’s health status), other studies show that there is a negativity bias when participants process self-related information, e.g., in the context of a performance situation^1,2,27,30–32^.

The present study aimed to test whether sleep plays a role in the retention, stabilization, revision and long-term maintenance of newly formed ability self-beliefs. To do so, we employed the Learning Of Own Performance (LOOP) task ^1,2,30^, in which participants form beliefs about their own abilities in novel domains based on manipulated social feedback. The task is particularly well suited for examining self-belief formation processes because feedback integration is not assessed explicitly through self-reports but instead captured implicitly through trial-by-trial belief updating in response to the manipulated and therefore targeted performance feedback^1,2^.

In a first belief formation phase participants were exposed to positive and negative feedback about their own cognitive estimation abilities with the belief formation being tested at the end of this phase (“Formation test”). During the subsequent 12-hour retention interval, participants either slept (at night; N=26) or remained awake (over daytime; N=28). The effects of sleep on the retention of formed self-beliefs were then assessed in a second test (“Post-retention test”). This phase was directly followed by a belief revision phase where participants again performed the LOOP task, but unbeknownst to them, were now provided with reversed feedback contingencies to their estimation abilities, thereby challenging previously experimentally established self-beliefs. After the revision phase, beliefs were assessed in a third test (“Revision test”). Finally, participants returned to the lab three weeks later for another assessment to probe the long-term maintenance of formed self-beliefs (“3-week test”). We hypothesized that sleep, compared to wakefulness, would facilitate the retention of the newly formed self-beliefs. Furthermore, we expected that the sleep-based consolidation would make self-beliefs more stable against their revision when exposed to conflicting feedback. We also hypothesized that these effects of sleep would persist at the 3-week follow-up test.

## Methods

### Participants

A total of 62 participants, aged between 18 and 28 years took part in a between-subjects design with participants assigned to either a Wake or Sleep group (see Figure 1A). Data from 8 participants were excluded from analyses (n=2 due to technical errors, n=1 due to insufficient SWS, n=5 due to a failure in adequately performing on the LOOP task, evidenced by a lack of variance or random fluctuations in their responses) leaving a final sample size of n=54, including n=28 Wake (mean age= 22.3, 18 female, 10 male) and n=26 Sleep participants (mean age= 22.0, 18 female, 8 male). There was further dropout of n=1 and n = 2 participants of the Sleep and Wake groups, respectively at the “3-week test”. A sensitivity power analysis conducted in G*Power 3.1 indicated that, with n=28 in the Wake and n=26 in the Sleep group, a two-sided α of .05 and 80% power, the minimum detectable between-group effect was Cohen’s d=0.78.

**Figure 1.**
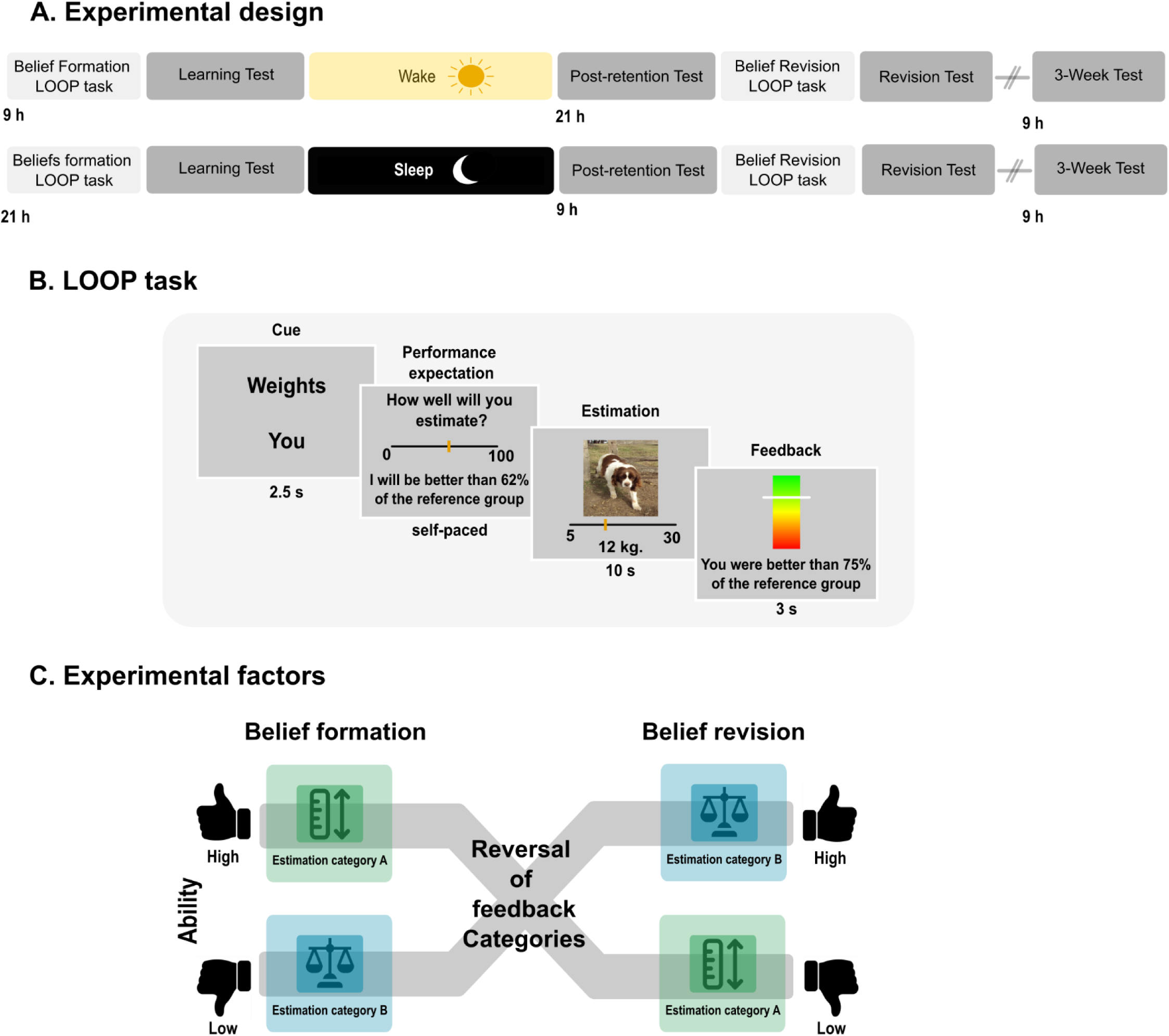
Experimental design and Learning Of Own Performance (LOOP) task (LOOP) **A. Experimental design**. The study consisted of a between-subjects repeated measures design. Both the wake and sleep groups started by performing the LOOP task either in the morning ∼ 9 h or evening ∼ 21 h. In both groups, the LOOP task was followed by a “Formation test” to assess a baseline measure of self-beliefs states before sleep/wakefulness. Next morning or evening, they performed the “Post-retention test”. Directly afterwards, subjects performed the LOOP task again but were now challenged by contradictory feedback. The LOOP task was followed by a “Revision test”. Three weeks later, all participants returned to the lab to perform a follow-up measurement “3-week test”. **B. LOOP task.** A first screen prompted the agent and the estimation category (e.g. weights). Participants then provided their performance expectation rating. Afterwards, they answered the actual estimation question and finally received feedback on their performance on that trial. This trial structure was repeated 10 times for each agent and estimation category (40 trials in total)**. C. Experimental factors.** The experimental factors of the LOOP task (High, Low) were assigned to one of the estimation categories (Estimation category A: height of buildings, B: distance between vehicles, C: weights of animals, D: number of objects), randomized and counterbalanced across participants. For the Belief Revision task, the estimation categories were reversed, meaning that those categories previously assigned for the high and low abilities, were interchanged. That way, participants faced feedback that contradicted their ability beliefs formed in the initial belief formation phase.

Participants were recruited at the University of Tübingen and provided written informed consent. Participants reported having a regular sleep-wake cycle and not carrying out shift work at least 6 weeks prior to the experiments. They did not present any history of neurological, psychiatric or endocrine disorder, did not take any medication at the time of the experiments, and were non-smokers. Participants of the Sleep group had an adaptation night in the sleep lab before the experiment proper. Participants received financial compensation for their participation. The study was approved by the ethics committee of the Medical Faculty of the University of Tübingen.

### Study design and general procedures

The experiment used a between-subjects design (Sleep vs. Wake) and comprised several phases (Figure 1A). It started with the belief formation phase where participants performed the LOOP task^1,2^ to form new ability self-beliefs based on social feedback, either in the evening (∼21:30 h; Sleep group) or in the morning (∼09:00 h; Wake group). The phase ended with the participants performing a “Formation test” assessing the initial level of the formed self-beliefs. Then, after self-belief formation, a 12-hour retention interval followed, which was filled with either sleep or wakefulness: The Sleep group spent one night in the sleep laboratory (including polysomnographic recordings between 23:00–07:00h), whereas the Wake group engaged in their usual daily activities (monitored by wrist actigraphy). After the retention interval, ability self-beliefs were again assessed (“Post-retention test”) and immediately followed by a belief revision phase, during which the participants performed the LOOP task again but now with reversed feedback contingencies (Figure 1C). The belief revision phase ended with the “Revision test” assessing changes in self-beliefs. Long-term maintenance of formed self-beliefs was assessed three weeks later in the “3-week test”.

### The Learning Of Own Performance (LOOP) task

The LOOP task allows participants to form novel beliefs about their own ability to estimate unfamiliar attributes or properties, such as the weight of animals or the height of houses (Figure 1B). The LOOP task has been implemented and extensively described in previous studies ^1,2,26,30–32^. While the task also includes a condition to examine how beliefs about another person’s performance are formed, this study concentrated on an assessment self-belief formation in the context of abilities. The assessment of the learning mechanisms underlying other-related belief formation was implemented only as a control condition (results of which are not reported here). To enable this control, the participant performed the LOOP task conjointly with another person (i.e., a confederate of the experimenter, sitting in an adjacent room), with the participants being told that they would take turns with the other person in either performing the estimation task themselves or estimating the other person’s performance.

Stimuli to be estimated were presented on a computer screen, and during each trial, participants received manipulated feedback about their assumed performance. There were four different categories for estimation (weight of animals, height of houses, number of objects, distance between vehicles), with two of these categories being randomly assigned to the *Self* and *Control* condition, respectively. One of the two categories was associated with overly high-performance feedback and the other with overly low-performance feedback on the assumed estimation ability. The respective mock feedback was based on the performance of a reference group of 350 participants tested before, and conveyed in terms of percentiles (e.g., "You are better than 65% of the reference group"). Feedback for the high estimation ability was normally distributed around the 65th percentile, feedback for the low estimation ability was normally distributed around the 35th percentile of the reference sample. High and low ability feedback was pseudo-randomly assigned to the estimated properties, and the categories were counterbalanced between *High vs. Low ability* and *Self* vs. *Control* conditions.

Each trial started with a note on the screen, indicating the type of task (*Self vs. Control*) and the type of property to be estimated (Figure 1B). Then participants were asked to indicate their expectation about how well they would perform (Performance expectation) on the same percentile scale that was used for performance feedback. Next, a picture of the respective property was shown, and participants were asked to estimate that property. Finally, they received the mock feedback about how well they had performed relative to the previously tested reference group (Feedback), before the next trial started. Overall, there were ten trials for each condition (Self/Control; High/Low ability) with feedback conditions randomized.

Participants were told they would receive additional money (i.e., up to 10 cents per trial) to increase adherence to the task. The intermediate money score was shown after approx. 10 trials to motivate honest answers and the best possible performance expectation. At the very end of the study, after three weeks, participants were informed about the controlled feedback structure of the task and received the full 5€ bonus.

### Revised Learning Of Own Performance task

The task structure was identical to that of the belief formation phase, except that the feedback contingencies for the *High vs. Low ability* categories were now - unknown to the participant - reversed. For example, if “estimating the heights of buildings” was the *Self-High* condition and “estimating the weights of animals” was the *Self-Low* condition during the LOOP, then these contingencies were switched in the belief revision phase (Figure 1C). In the belief revision phase, also pictures with new animals and houses were used.

### Testing beliefs

Belief states were tested at four different time points: “Formation test”, “Post-retention test”, “Revision test”, and “3-week test”. (Figure 1A). All tests followed the same procedure, i.e., participants completed 20 trials (five per feedback condition) in which they estimated the properties of novel images using the same procedure as in the LOOP task. In addition to each performance expectation, participants were asked to indicate their confidence in the expected performance (e.g., "How sure are you that you will do better than x%?"; see Supplementary Material). Importantly, no feedback on estimation performance was provided, to avoid new belief formation processes.

### Debriefing

In the very end of the study, participants underwent an interview where they were debriefed about the cover story. They were told that their computer was not connected to that of the other participant during the estimation task and that they had received manipulated feedback. They were also informed that the main variables of the study were not the estimation performance per se, but their change in expectation during the course of the experiment.

### Control variables and psychometric tests

After the experiment proper, participants completed the Self-Description Questionnaire (SDQ), Beck depression questionnaire (BDI-II), Social Interaction Anxiety Scale (SIAS), and the Social Phobia Scale (SPS). Participants also completed a vigilance task before the experiment proper and the Stanford Sleepiness Scale (SSS). Additionally, participants of the Sleep group filled out a questionnaire on sleep quality in the morning after the experimental night, and participants of the Wake group a questionnaire on their daily activities during the daytime retention interval.

### Sleep recording

Polysomnography was continuously recorded with a BrainAmp MR plus amplifier system (Brain Products GmbH, Germany) with scalp electroencephalogram (EEG) electrodes positioned according to the international 10–20 system at C3 and C4, referenced to electrodes on the mastoids. One ground electrode was placed on the forehead. Impedances were kept below 5 kΩ. EEG data were sampled at 200 Hz and saved on a computer for offline analyses. In addition to the above setup, two electrodes were attached to record horizontal and vertical electrooculography, and two electrodes were attached on the chin to record electromyography. Visual scoring of 30-second polysomnographic recordings followed standard scoring criteria ^33^. Sleep stages S1, S2, SWS (S3 + S4), REM sleep, wake, epochs containing movement as well as other arousals were identified from lights off until participants woke up. Slow oscillations and sleep spindles detection were performed using the open-source toolbox SpiSOP (https://www.spisop.org, RRID:SCR_015673) (see Supplemental Methods for details).

### Actigraphy

Participants of the Wake group were continuously monitored during the retention interval by means of an Actiwatch 2 (Philips Respironics, Amsterdam, Netherlands) in order to ensure that they did not sleep.

### Statistics

Statistical analyses for behavioral and sleep data were performed in Jamovi 2.6.44.0 and Matlab R2024b.

The belief formation process during the LOOP and revised LOOP was quantified by first calculating the change in performance expectations between consecutive trials within the same ability condition on a trial-by-trial basis resulting in trial-wise update scores. Updates were then averaged within each experimental condition and further analyzed with mixed analyses of variance (ANOVAs) comprising a within-subjects factor “Ability”, representing the high vs low ability feedback conditions, and a group factor “Sleep/Wake”. This procedure differed from our previous publications^2,22^ in which we typically estimated Rescorla-Wagner learning models. The change was motivated by the reduced number of trials per condition and the altered feedback structure in the present study as we used fixed feedback sequences (as described above) rather than predetermined prediction-error sequences^1,2,21,22,26^.

For each of the four tests of belief states (i.e. “Formation test”, “Post-retention test”, “Revision test”, “3-week test”), performance expectation was calculated separately for the two feedback conditions (High and Low). The current belief state during the test was determined by the mean performance expectations across all five trials of each ability condition. Statistical comparisons were done with mixed analyses of variance (ANOVA) comprising a within-subjects factor “Ability”, representing the high vs low ability feedback conditions, and a group factor “Sleep/Wake”. In case of significant ANOVA interaction effects, post-hoc t-tests were calculated and Bonferroni correction for multiple comparisons was performed if indicated. Exploratory correlations between measures of self-belief formation and sleep parameters were conducted using Spearman correlation coefficients and are reported uncorrected for multiple testing. A value of P<0.05 was considered significant. In the absence of significant differences between conditions, Bayesian repeated-measures ANOVA–based model comparisons were conducted using default Cauchy priors on fixed effects (r = 0.5).

## Results

### Successful Self-Belief Formation during the LOOP Task

We first verified successful formation of new ability self-beliefs on the LOOP task in both the Sleep and Wake groups. To do so, self-belief formation was quantified on a trial-by-trial basis by calculating the change in expected performance between consecutive trials within the same ability condition (Figure 1; see Methods). Participants’ initial performance expectations, taken from the first trial during the belief formation phase, scattered around the center of the distribution and did not differ between the groups (Wake: mean = 49.8; Sleep: mean = 52.4; main effect Sleep/Wake: *F_1, 52_* = 1.74, *p* = .193, part. η^2^ = 0.03; main effect Ability: *F_1, 52_* = 0.28, *p* = .601, part. η^2^ = 0.005; interaction: *F_1, 52_* = 0.08, *p* = .784, part. η^2^ = 0.001; mixed ANOVA). This allowed us to steer the participants into both ability self-belief directions by giving them either overly high or overly low feedback on their assumed estimation performances for each condition, respectively.

Over the course of the belief formation phase, participants in both groups used the manipulated feedback to update their performance expectations towards lower and higher estimation ability self-beliefs. This was indicated by a significant difference in the mean updates of performance expectations between the low and high ability feedback condition (Ability main effect: *F_1, 52_* = 171.60, *p* < .001, part. η^2^ = 0.77; Figure 2A). The analysis did neither reveal a significant difference between groups (main effect Group: *F_1, 52_* = 0.52, *p* = .476, part. η^2^ = 0.010) nor a Group x Ability interaction (*F_1, 52_* = 1.90, *p* = .174, part. η^2^ = 0.035), indicating that both groups similarly updated their performance expectations in response to the feedback and arrived at comparable levels of ability self-beliefs at the end of the belief formation phase. The “Formation test” confirmed a significant effect of ability (*F_1, 52_* = 251.83, *p* < 0.001), no significant Group x Ability interaction (*F_1, 52_* = 0.48, *p*=0.49), nor a main effect of Group *(F_1, 52_* = 2.69, *p*=0.107).

**Figure 2.**
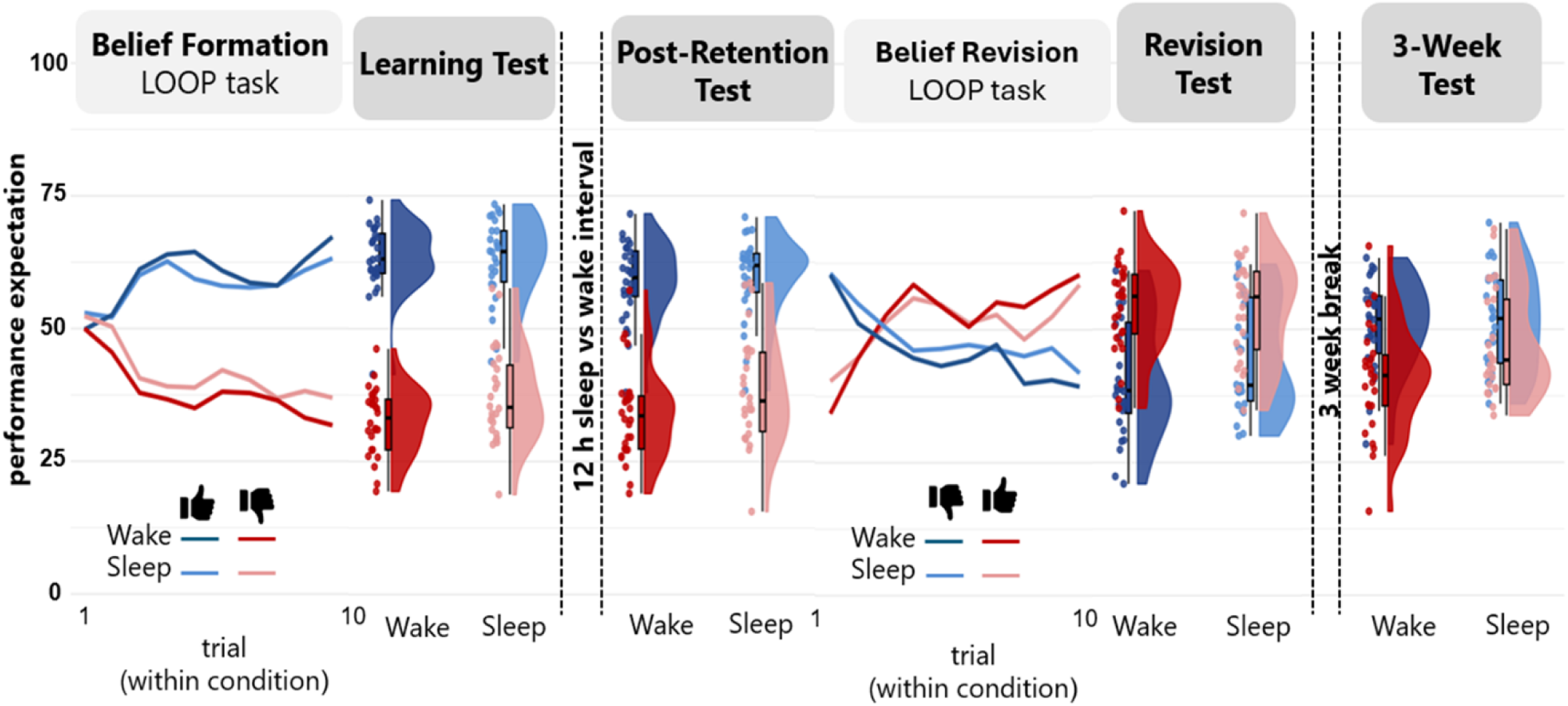
Self-belief formation, belief revision, and four state assessments. The study followed a between-subjects repeated-measures design, with participants assigned to either a sleep or wake group and encountered high and low mocked feedback. Initial performance expectation scattered around the center of the distribution, confirming there were no initial differences between the wake and sleep groups. Belief formation started with the LOOP task, where participants encountered either high (shown in blue) or low feedback on their estimation performances (shown in red), both in the wake (dashed lines) and sleep group (full lines). Immediately after the LOOP task, a baseline belief state measurement (“Formation test”) was conducted to assess self-beliefs prior to the sleep/wake retention interval. After the 12 h sleep/wake retention interval, belief states were assessed again in the “Post-retention test”, confirming successful belief formation ∼12 hours before, yet no immediate effect of sleep vs. wakefulness on belief states post-consolidation. Next, participants performed the LOOP task again but now with reversed feedback contingencies. Results showed that participants adjusted their self-beliefs according to the new feedback, indicating successful belief revision as confirmed in the “Revision test”. After three weeks, assessment of belief states suggested a more positive memory for the initially formed low ability category in the sleep compared to the wake group.

### Sleep did not Affect the Retention of Formed Self-Beliefs

To assess whether Sleep vs. Wake were related to the retention of newly formed ability self-beliefs, we compared self-belief states between the “Formation test” and the “Post-retention test” (using an additional Pre/Post ANOVA factor). Sleep did not affect the retention of formed ability self-beliefs after 12h when compared to the Wake group (*F_1, 52_* = 0.24, *p* = .630, part. η^2^ = 0.004, for respective Sleep/Wake x Ability x Pre/Post interaction). Sleep did also not change the overall positivity or negativity of beliefs (Sleep/Wake x Pre/Post: *F_1, 52_* = 0.22, *p* = .644, part. η^2^ = 0.004). The Sleep/Wake main effect and Sleep/Wake x Ability interaction were likewise non-significant (*p* > .54). Bayesian repeated-measures ANOVA-based model comparison with default Cauchy priors on fixed effects (r = 0.5) showed that the model including Group (sleep vs. wake) and its interactions was strongly disfavored relative to the corresponding model without Group (BF₁₀ = 0.013; see Table S4). This indicates strong evidence against a sleep-related effect on the retention of newly formed ability self-beliefs.

### No Effect of Sleep on Belief Revision

In the belief revision phase, participants again performed the same estimation tasks but on different stimuli (see Methods). Unbeknownst to them, the previously high-ability category was now paired with low-ability feedback, and vice versa (Figure 1C). Belief updates for the belief revision phase were calculated in the same way as for the belief formation phase, i.e. as the change in expected performance between consecutive trials within the same ability condition based on participants’ trial-by-trial performance expectations, and mean updates within conditions were used for further analyses. Participants in both groups revised their initially formed beliefs as confirmed by a significant main effect of Ability in an analysis on the mean updates (*F_1, 57_* = 186.31, *p* < .001, part. η^2^ = 0.78). As in the belief formation phase, Sleep and Wake groups did not differ in their performance during the belief revision phase (*F_1, 52_* = 3.57, *p* = .064, part. η^2^ = 0.064, for Sleep/Wake main effect; *F_1, 52_* = 1.47, *p* = .231, part. η^2^ = 0.027, for Sleep/Wake x Ability interaction).

To additionally assess whether prior sleep in the Sleep group stabilized newly formed ability self-beliefs, we compared the updating between the belief formation phase with the belief revision phase using an additional “Belief formation/Belief revision” ANOVA factor. This analysis did not reveal any significance for the respective Sleep/Wake x Ability x Belief formation/Belief revision interaction (*F*(1,52) = 0.03, p = .868, η² = .001), indicating that the revision of self-beliefs operated similarly to how self-beliefs were formed in the first place and did not differ between Sleep and Wake groups. Sleep, likewise, did not alter the overall positivity or negativity of self-beliefs across conditions updates (Sleep/Wake x Belief formation/Belief Revision: *F_1, 52_* = 0.46, *p* = .501, part. η^2^ = 0.009). Also, the Sleep/Wake main effect (*p* = .064) and Sleep/Wake x Ability interaction (*p* = .129) were not significant. Bayesian repeated-measures ANOVA–based model comparison with default Cauchy priors on fixed effects (r = 0.5) showed that the model including Group (sleep vs. wake) and its interactions was strongly disfavored relative to the corresponding model without Group (BF₁₀ = 0.033; see Table S5). This indicates strong evidence against a sleep-related effect on self-belief retention, which would have been signified by less revision of, i.e. more adherence to, a-priorily formed ability self-beliefs.

The comparison between the “Post-retention test” and the “Revision test” confirmed a significant main effect of Ability (*F_1, 52_* = 20.59, *p*<0.001) no significant three-way Sleep/Wake x Ability x Post-Retention test/Revision test interaction (*F_1, 52_* =0.85, *p*=0.35) nor a main Group effect (*F_1, 52_* =2.43, *p*=0.125).

### Positivity Bias after 3-Weeks in the Sleep Group

To assess potential long-term effects of sleep, participants completed a follow-up test three weeks later. Sleep specifically affected the long-term maintenance of initially formed low-ability beliefs, with participants who slept recalling them as more positive at the “3-week test” than the wake group participants (F_1, 49_ = 6.60, p = .013, part. η^2^ = 0.119, for respective Sleep/Wake x Ability x Belief revision/3-week test interaction, *t_49_* = 2.06, *p* = .044, Cohen’s d = 0.58, for respective post hoc pairwise comparison between Sleep and Wake group, Fig. 3A). A supplementary comparison on all four tests (“Formation test”, “Post-retention test”, “Revision test” and “3-week test”) revealed that the sleep group’s “3-week test” was the only test condition without significant differences between initially formed low and high ability self-beliefs (Fig. 3A). This pattern may suggest that the sleep group returned to their initial self-beliefs prior to the belief formation phase, potentially indicative of increased forgetting as compared to the Wake group.

**Figure 3.**
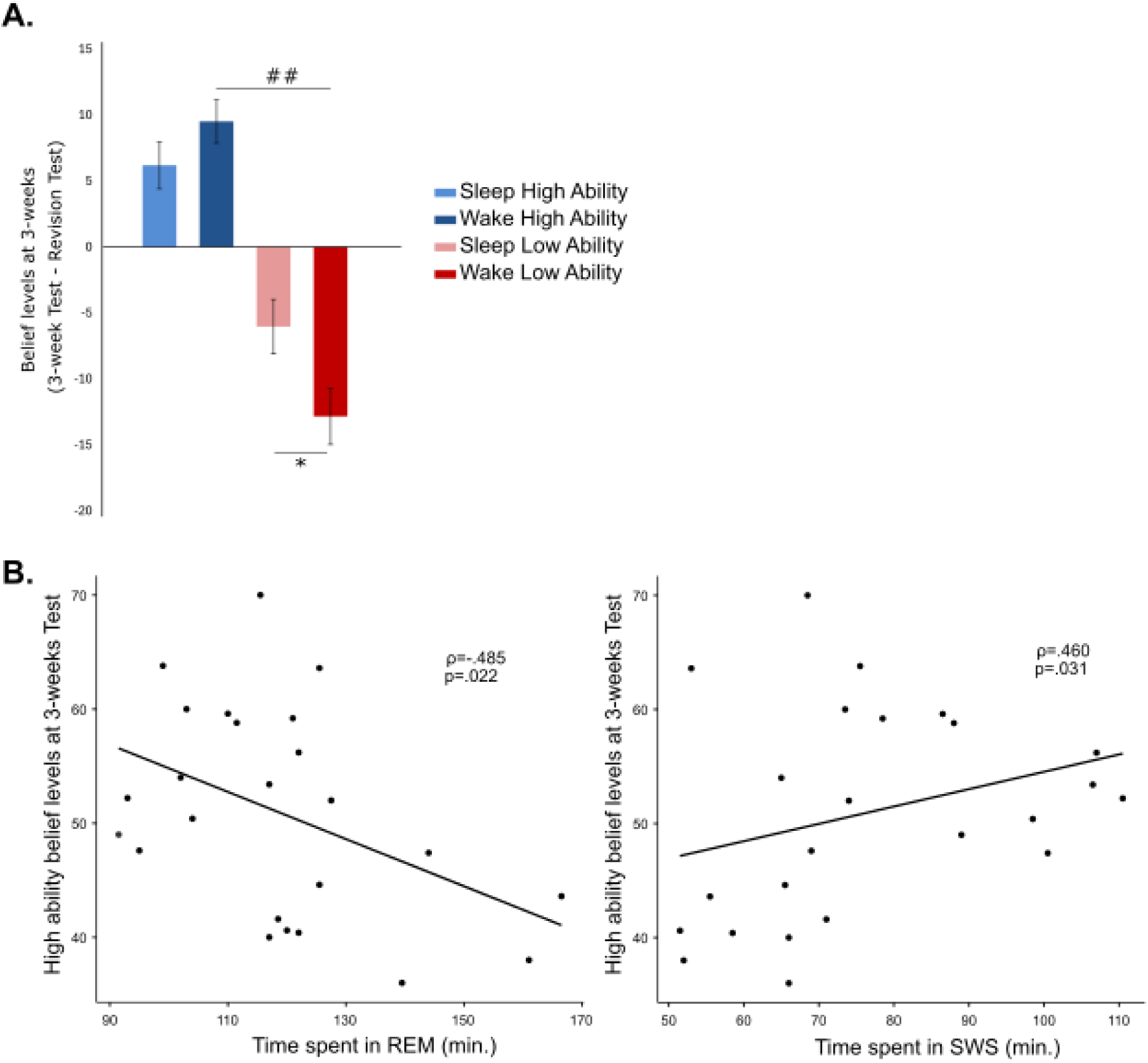
A. Belief levels at the “3-week test” following the belief revision phase (3-week test - Revision test) for High and Low ability self-beliefs in the Sleep and Wake groups. Low ability self-beliefs in the Sleep group showed a shift toward more positive expectations compared with the Wake group. Bars represent mean ± SEM. * p < 0.05, for pairwise comparisons between Sleep and Wake groups. ## p < 0.01, for pairwise comparison with low ability self-belief for respective group. 3B. Time spent in REM sleep and high-ability beliefs after three weeks were negatively correlated (Spearman’s ρ = −.485, p = .022), and time spent in SWS sleep and high-ability beliefs after three weeks were positively correlated (Spearman’s ρ = .460, p = .031).

### Interindividual Differences in Sleep Parameters and Belief Expectations

To examine whether sleep following self-belief formation was linked to subsequent belief states, we conducted correlation analyses between belief states at the “Post-retention test” session and sleep parameters, including time spent in each sleep stage, slow oscillation count, and fast and slow spindle count. These analyses were controlled for BDI scores, as depressive symptoms can influence both sleep quality^34^ and self- beliefs^31^, as well as for initial self-beliefs (as assessed by first trial of the belief formation phase).

At the uncorrected level, a negative association emerged between REM sleep duration and belief states in the high ability condition at the “3-week test” (Spearman’s ρ = −.485, p = .022), together with a positive association between SWS duration and belief states in the high ability condition (Spearman’s ρ = .460, p = .031), suggesting that these sleep stages may differentially contribute to the long-term maintenance of newly formed ability self-beliefs (Figure 3B). It is important to notice that one participant that exhibited an elevated SWS value relative to the sample distribution was excluded from the correlation analyses, however when including this subject both correlations remained significant (REM: ρ = −.532, p = .009; SWS: ρ = .507, p = .014). No other correlations reached significance. Given the number of tests conducted, these findings should nevertheless be interpreted with caution.

## Discussion

Here, we investigated for the first time whether sleep facilitates the retention, stabilization, revision and long-term maintenance of newly formed ability self-beliefs. We replicated successful self-belief formation using the LOOP task^1,2,22,26^. However, nocturnal sleep after belief formation did not enhance the retention of these newly formed ability self-beliefs in comparison with a daytime wake control group. Sleep also did not affect subsequent revision of these newly formed self-beliefs, when participants performed the LOOP task a second time but now with switched feedback contingencies. This indicates that sleep did not have a stabilizing effect on newly formed ability self-beliefs as otherwise self-beliefs would have been less prone to their revision. After three weeks, ability self-beliefs in the sleep group returned to the a-priori level with a tendency towards a shift of newly formed low-ability self-beliefs (i.e. “I am not good at…”) into a more positive direction. This pattern of results contradicts our initial hypothesis that sleep would strengthen experimentally formed self-beliefs and thereby contribute to the consolidation of self-beliefs, which are central to human social functioning.

First, our experiments replicate previous studies using the LOOP task, allowing participants to successfully form^1,2,22,26^ and revise self-beliefs in response to self-related performance feedback, independently of the sleep or wake condition. Although research on the link between sleep and the formation of self-beliefs is scarce, related work suggests that sleep can modulate social information processing. Studies using targeted memory reactivation (TMR) suggest that social feedback and social evaluations can be selectively reactivated during sleep^19,35^. Complementary evidence comes from sleep deprivation studies, showing that reduced sleep increases susceptibility to changes in beliefs ^36^. Likewise, sleep after a learning intervention on gender and race stereotypes was shown to robustly strengthen newly learned concepts ^15–19^. Against this backdrop the failure of sleep to strengthen newly formed ability self-beliefs in the present study is surprising, but might be related to the fact that this study – unlike those previous studies – concentrated on self-related learning processes ^1,2^. Compared with the formation of beliefs about another, typically unfamiliar person, the formation of new self-beliefs may qualitatively differ, as the representation of self-beliefs is anchored in rather overlearned and enduring schema-like memories^21^. Thus, any new social feedback information related to the self needs to be assimilated to these pre-existing and therefore quite precise self-beliefs (or schema-like memories) ^2^. Feedback information conflicting with such pre-existing self-beliefs may even be aborted from the assimilation process preventing premature consolidation of such beliefs during sleep. Indeed, findings with non-social learning materials suggest that sleep preferentially strengthens memories conformant with pre-existing information or concepts ^11,37,38^. Accordingly, the failure of sleep to strengthen newly formed self-beliefs in the present study could similarly reflect the strongly enhanced difficulty to assimilate new self-related information into pre-existing self-beliefs.

Alternatively, the absence of a sleep-related strengthening of newly formed self-beliefs may be attributed to time-of-day influences rather than sleep-dependent consolidation processes. Previous work using the LOOP task found that participants tested in the evening had higher learning rates than those tested earlier in the day, and that these differences varied rhythmically across the day^39^. These findings suggest that self-belief formation processes may be influenced by time-of-day effects and circadian rhythms, although the precise mechanisms involved, such as chronotype, affective state, attention, or prior sleep, remain to be elucidated ^39^. In the present study, the LOOP task was conducted in the evening for the sleep group and in the morning for the wake group. Although belief states after the belief formation phase were comparable between the groups, underlying circadian confounds impacting the belief formation phase may have blurred the effects of subsequent sleep.

Despite the absence of sleep-related effects on short-term belief retention and revision, interesting findings emerged at the three-week follow-up. Specifically, participants who slept following the belief formation phase showed a selective shift toward more positive recall of initially formed low ability self-beliefs. Previous research using the LOOP task has demonstrated that the formation of beliefs about one’s own ability is strongly influenced by negative feedback when it is perceived as informative or useful for improvement, with participants incorporating negative feedback more strongly than positive feedback ^1,2,30^. Against this background, the present findings on the three-week effect suggest that although negative feedback can initially drive belief updating^31^, such effects may weaken over time, particularly following direct sleep after learning. Notably, prior work using the LOOP task has primarily focused on short-term effects of belief formation and revision typically occurring within several hours ^1,2,30^. To date, no studies have examined dynamics of self-belief formation over longer retention intervals, making the present three-week follow-up a novel contribution to understanding long-term maintenance of newly formed self-beliefs.

Another reasoning about this finding could be that sleep not only failed to strengthen the newly formed self-beliefs but kept them in a more labile state. Unlike the wake group, participants in the sleep group gradually attenuated and may have forgotten the more negative self-related feedback information as processed during the belief formation phase. This pattern would be consistent with proposals that sleep can also promote selective forgetting, e.g., by favoring the consolidation of schema-like representations over context-specific details^40,41^. Indeed, the long-term changes in self-beliefs observed after sleep might reflect a competition between two belief representations: a newly acquired, LOOP task-induced ability self-belief, and a pre-existing, more stable trait-like global self-belief held prior to the experiment that is strengthened by sleep. In this line, the outcome of the sleep-dependent consolidation process would be determined by the conformity of the newly processed information with pre-existing self-beliefs^8^. Notably, this interpretation is also partly supported by our exploratory sleep analyses. Greater time spent in SWS was associated with stronger maintenance of high ability self-beliefs at the three-week follow-up, whereas greater REM sleep was associated with weaker maintenance. Although exploratory, these findings are consistent with previous work assigning complementary roles to SWS and REM sleep, with SWS supporting the stabilization of recently acquired information and REM facilitating their subsequent transformation or integration with pre-existing knowledge ^8,11^.

## Conclusion

In conclusion, against our hypotheses, the present findings failed to show an enhancing effect of sleep on the retention, stabilization, revision and long-term maintenance of newly formed ability self-beliefs. Instead, sleep appeared to even weaken the initially induced low-ability self-beliefs over extended time scales leading to a more positive recall. Further research is warranted to scrutinize the robustness of the facilitating effect of a positive long-term shift in self-belief formation after sleep and its underlying mechanisms.

## Author contributions

CB and SD planned and designed the study set-up. SK, AS, NC and LMP developed the LOOP task and contributed to the set-up of the study. LE collected the data. JC, JB, SK and LMP analyzed the data. JC, SK and LMP wrote the first draft of the manuscript. All authors discussed the data and revised the manuscript.

## Data availability

Data will be made available upon reasonable request.

