## Supplementary materials for "The role of sleep in the retention and long-term maintenance of newly formed ability self-beliefs"

**Supplementary methods**

**Sleep spindle detection**

Sleep spindle detection was performed using the open-source toolbox SpiSOP ([https://www.spisop.org](https://www.spisop.org/) , RRID:SCR_015673) implemented in MATLAB (MathWorks, Natick, MA) and used in several previous studies^1,2^. Based on the individual power spectra during sleep stages S2, S3 and S4, peak frequencies for slow and fast spindles were determined separately for each participant. Detection was then performed individually for each subject and electrode using a 2-Hz frequency band centered on the respective peak frequency (±1 Hz). For each participant, the root mean square (RMS) signal was calculated using a 0.2 sec window, smoothed by a moving average (0.2 sec window). A  spindle was detected whenever this smoothed moving RMS window exceeded an individual threshold (1.5 standard deviations of the filtered signal in the respective channel) for 0.5–3 s. Slow and fast spindle counts were calculated for each participant and averaged across C3 and C4 electrodes.

**Slow Oscillation detection**

Slow Oscillations (SOs) detection during non-REM sleep epochs was also performed using SpiSOP. The EEG signal was bandpass filtered between 0.3 and 4 Hz. Time intervals between consecutive positive-to-negative zero crossings corresponding to frequencies between 0.5 and 1.25 Hz (0.8–2 s) were identified as potential SOs. For each channel, the mean negative peak potential and the mean peak-to-trough amplitude were calculated across all potential events. Only those events were considered as SOs whose negative peak potential was lower than 1.25 times the mean negative peak potential and whose peak-to-trough amplitude exceeded 1.25 times the mean amplitude of all putative SOs. SO count was calculated for each participant separately for C3 and C4 and averaged across electrodes.

**Certainty Ratings**

A mixed ANOVA on certainty ratings including all test time points revealed no main effect of Group, *F*(1,49) = 0.68, p = .41, and no significant interactions involving Group (all p ≥ .210), indicating that certainty ratings did not differ between Sleep and Wake conditions. However, a significant main effect of Time was observed, *F*(3,147) = 14.54, p < .001, reflecting a general decrease in certainty across time points (See Table S3, supplementary materials)

**Supplementary Tables**

**Table S1.** Expectation performance (mean ± SEM)

| **Test session** | **Sleep (Mean ± SEM)** | **Wake (Mean ± SEM)** |
| --- | --- | --- |
| High Ability Learning Test | 62.9 ± 1.54 | 61.7 ± 1.58 |
| Low Ability Learning Test | 37.4 ± 1.75 | 33.9 ± 1.39 |
| High Ability Post-Retention Test | 59.7 ± 1.51 | 57.5 ± 1.61 |
| Low Ability Post-Retention Test | 38.6 ± 2.07 | 34.9 ± 1.70 |
| High Ability Revision Test | 44.7 ± 2.07 | 40.8 ± 2.12 |
| Low Ability Revision Test | 53.9 ± 1.85 | 53.6 ± 1.59 |
| High Ability 3-Week Test | 51.5 ± 1.89 | 50.5 ± 1.71 |
| Low Ability 3-Week Test | 47.4 ± 2.02 | 41.4 ± 2.14 |

n: Sleep = 26 (3-Week Test=25), Wake = 28 (3-Week Test =26).

**Table S2.** Mean-trial-by-trial updates (mean ± SEM)

| **LOOP task session** | | **Sleep (Mean ± SEM)** | **Wake (Mean ± SEM)** |
| --- | --- | --- | --- |
| Belief formation high ability | 1.24 ± 0.268 | | 1.75 ± 0.221 |
| Belief formation low ability | −1.69 ± 0.260 | | −1.86 ± 0.229 |
| Belief revision high ability | 2.01 ± 0.283 | | 2.81 ± 0.272 |
| Belief revision low ability | −2.09 ± 0.287 | | −2.09 ± 0.264 |

n: Sleep = 26, Wake = 28.

**Table S3.** Certainty ratings (mean ± SEM)

**Test session	Sleep (Mean ± SEM)	Wake (Mean ± SEM)**

High Ability Learning Test	59.8 ± 1.93	58.9 ± 2.77

Low Ability Learning Test	62.0 ± 2.37	60.4 ± 2.33

High Ability Post-Retention Test	58.4 ± 2.23	54.2 ± 2.90

Low Ability Post-Retention Test	59.4 ± 2.80	57.1 ± 2.33

High Ability Revision Test	53.4 ± 2.82	52.4 ± 3.73

Low Ability Revision Test	53.1 ± 2.34	48.5 ± 3.03

High Ability 3-Week Test	51.1 ± 3.43	47.1 ± 3.48

Low Ability 3-Week Test	51.3 ± 3.18	51.9 ± 2.92

n: Sleep = 26 (3-Week Test=25), Wake = 28 (3-Week Test =26).

**Table S4. Bayesian statistics.** Results section 2. Sleep did not affect belief retention after a 12 hs interval. Bayesian repeated measures ANOVA with within factors ability (Low vs High), time (Learning Test vs Post-retention Test) and between subjects factor group (Sleep vs Wake)

| Model Comparison | | | | | |
| --- | --- | --- | --- | --- | --- |
| **Models** | **P(M)** | **P(M\|data)** | **BF_M_** | **BF_10_** | **error %** |
| Null model (incl. Abilty, Time, Abilty✻ Time, subject) | 0.167 | 0.44484 | 4.0064 | 1.0000 |  |
| group | 0.167 | 0.34253 | 2.6050 | 0.7700 | 3.97 |
| group + group✻ Abilty | 0.167 | 0.10530 | 0.5884 | 0.2367 | 4.68 |
| group + group✻ Time | 0.167 | 0.07296 | 0.3935 | 0.1640 | 5.85 |
| group + group✻ Abilty + group ✻ Time | 0.167 | 0.02856 | 0.1470 | 0.0642 | 26.74 |
| group + group✻ Abilty + group ✻ Time + group ✻ Abilty ✻ Time | 0.167 | 0.00581 | 0.0292 | 0.0131 | 6.07 |
| Note. All models include Abilty, Time, Abilty✻ Time, subject. | | | | | |

**Table S5. Bayesian statistics.** Results section 3. Sleep did not affect belief revision. Bayesian repeated measures ANOVA on the mean trial-by-trial updates with within factors LOOP (Formation vs Revision) x Ability (Low vs High) and between subjects factor group (Sleep vs Wake).

| Model Comparison | | | | | |
| --- | --- | --- | --- | --- | --- |
| **Models** | **P(M)** | **P(M\|data)** | **BF_M_** | **BF_10_** | **error %** |
| Null model (incl. Ability, Time, Ability✻ Time, subject) | 0.167 | 0.4492 | 4.0772 | 1.0000 |  |
| group | 0.167 | 0.1997 | 1.2476 | 0.4446 | 13.69 |
| group + group✻ Ability | 0.167 | 0.2412 | 1.5894 | 0.5370 | 7.63 |
| group + group✻ Time | 0.167 | 0.0371 | 0.1925 | 0.0825 | 6.24 |
| group + group✻ Ability + group ✻ Time | 0.167 | 0.0580 | 0.3080 | 0.1292 | 7.79 |
| group + group✻ Ability + group ✻ Time + group ✻ Ability ✻ Time | 0.167 | 0.0148 | 0.0753 | 0.0330 | 6.66 |
| Note. All models include Ability, Time, Ability✻ Time, subject. | | | | | |

**Table S6. Sleep scoring and oscillations**

| **Sleep Variable** | **Mean ± SEM** | **Unit** |
| --- | --- | --- |
| WASO (Wake After Sleep Onset) | 10.6 ± 2.09 | min. |
| Stage 1 (S1) | 46.1 ± 3.39 | min. |
| Stage 2 (S2) | 238 ± 3.93 | min. |
| Stage 3 (S3) | 51.3 ± 3.03 | min. |
| Stage 4 (S4) | 30.6 ± 4.43 | min. |
| Slow Wave Sleep (SWS) | 81.8 ± 5.86 | min. |
| REM Sleep | 115 ± 4.45 | min. |
| Sleep Duration (onset to offset) | 497 ± 2.49 | min. |
| Total Sleep Time (TST) | 481 ± 3.56 | min. |
| Fast Spindle Count (12-15 Hz) | 1268 ± 112 | - |
| Slow Spindle Count (9-12 Hz) | 796 ± 83.3 | - |
| Slow Oscillation Count (0.5-1 Hz) | 1137 ± 55.4 | - |

n: Sleep = 26
